# Quantifying direct and indirect maternal effects on male vervets’ rank post-dispersal

**DOI:** 10.64898/2026.10.29.762785

**Authors:** Alba Motes-Rodrigo, Pooja Dongre, Varun Manavazhi, Erica van de Waal

## Abstract

Early life experiences have important fitness consequences across animals, with maternal environments playing a crucial role in some species’ survival and reproduction during adulthood. A potential pathway linking maternal behaviour with individual fitness is dominance rank, which in some primates – like cercopithecines – is maternally inherited during the juvenile period and strongly linked to individual social behaviour and reproductive success. While females of these species maintain their rank for life in their natal group, males disperse and must integrate into new dominance hierarchies. In this context, the long-term effects of maternal characteristics on male rank post-dispersal and the potential of males’ social behaviour in mediating this relationship remains unexplored. Using 16 years of data from a wild vervet monkey population (*Chlorocebus pygerythrus*), we tested the effects of maternal rank, grooming and proximity on the rank males acquired their first year post-dispersal. In addition, we measured the extent to which the males’ own social behaviour both pre– and post-dispersal mediate this relationship combining Bayesian social networks with mediation analyses. We did not find support for a mediating effect of males’ social behaviour on the relationship between maternal characteristics and male rank post-dispersal. Instead, we found that maternal characteristics influence males’ social behaviour during the juvenile period and that males’ own social behaviour post-dispersal (grooming out-degree) is related to their rank when they integrate into a new group. In addition, we found that consistency in males’ social behaviour pre-to post-dispersal is restricted to grooming behaviour, highlighting the different roles of proximity and grooming relationships in vervet societies. Taken together these results reveal very limited long-term effects of maternal characteristics on male behaviour during the heavily underexplored process of dispersal and the flexibility of male vervet social behaviour following a change in social environment.

## Introduction

The quality of the social environment an individual experiences has important consequences for survival and reproductive success across taxa^1^. Social environment quality can be measured based on various components that not only directly contribute, but also interact, in shaping Darwinian fitness. These include social integration, social status and the degree of adversity experienced in early life. Previous studies both in humans and non-human animals have shown that better integrated individuals have healthier and longer lives^2^. Indeed, social isolation in humans increases the risk of cardiovascular disease by 30%^3^, the risk of premature dementia by 50%^4^ and overall mortality risk by 29%^5^. Recent meta-analyses have found strikingly similar patterns to those found in humans across primates, equids, cetaceans and rodents, where social integration increases the odds of survival by 20 to 70%^6^.

Social status in group-living animals that often feed on monopolizable resources is generally determined by their position in the groups’ hierarchy^7^. These positions are represented by dominance ranks which are usually based on an individual competitive ability and are estimated based on conflicts’ wins and losses^8^. Dominance rank is positively associated with access to resources and mates, health^9^, reproductive success, and lifespan^10,11^ For instance, in spotted hyenas (*Crocuta crocuta*), higher-ranking females reach reproductive maturity earlier, have shorter inter-birth intervals and produce more surviving offspring than lower-ranking females^12^. Similarly, higher ranking females across primate species have shorter inter-birth intervals (e.g., vervet monkeys, *Chlorocebus pygerythrus*^13^) and higher infant survival (e.g., vervets^13^; chimpanzees, *Pan troglodytes*^14^). In turn, higher ranking males in many primate species have higher fecundity and mating success than subordinate males^10^.

An individual’s rank and its social behaviour are often interlinked. Across many primates, higher ranking individuals generally receive more grooming from more conspecifics and are frequently found in close proximity to other group members^9,15^. However, the exact relationship between social behaviour and rank can vary among species and populations. In chimpanzees, alpha males in some populations have the highest number of grooming partners and grooming rates^16^ whereas in other populations, rank and grooming degree are unrelated^17^. In captive rhesus macaques (*Macaca mulatta*), higher ranking individuals receive more grooming^18,19^ and in Bonnet macaques (*Macaca radiata*) high-ranking males and females are found more often in close proximity to conspecifics^20^. In addition to being a correlate of rank, social behaviour can also act as a driver. For example, grooming socially central females and spending time in their close proximity was found to lead to higher ranks in vervet monkeys^21^.

Regarding early-life adversity, it is well established that experiencing disadvantageous conditions during development affects fitness outcomes later in life including reproductive timing, disease risk and premature death^22^. Conversely, in non-human mammals and birds, higher environmental quality during development measured in terms of food availability, lack of climatic hardship and lower population density, has been linked to slower rates of reproductive senescence^1^. In those species with extended dependent periods such as primates, an important component of early life environments is maternal sociobehaviour^22^. Mothers are a fundamental source of early-life support and both their social status as well as investment in their offspring can impact fitness outcomes beyond their offspring’s dependent period. Maternal rank regulates offspring access to influential partners as well as resources such as water^23^ and food^24^ which results in better body condition during juvenescence. This preferential access to resources, often leads immatures born of higher-ranking mothers to wean sooner without compromising survival^25^ as well as to have higher reproductive fitness (“silver-spoon effect”^26^). For example, daughters of higher-ranking mothers reach sexual maturity earlier than daughters of lower-ranking mothers in chimpanzees^14^ and give birth earlier to their first litter in spotted hyenas^27^.

Critically, maternal rank often influences offspring rank. The process by which offspring obtains ranks adjacent to their mothers is known as rank inheritance^28–31^ and it can operate via different pathways. For instance, directly inheriting traits linked to competitive ability such as body size or aggressiveness might predispose offspring to hold similar ranks as their mothers^30^. Maternal rank might also affect offspring rank via exposure to rank-related hormones during pregnancy that influence their competitive ability. In baboons (*Papio cynocephalus*), subadult male glucocorticoid levels were mainly predicted by the mother’s rank at the time of conception rather than individuals’ own age and rank^32^, with sons of low-ranking mothers having higher levels than sons of high-ranking females. Similarly, in hyenas, offspring of higher-ranking females are exposed to higher androgen levels in the uterus which has been related to higher aggression levels during the juvenile period^33^. Offspring rank can also be modulated by maternal behaviour postpartum^34^. In vervet monkeys for instance, the initial ranks of sons and daughters of subordinate mothers is established around five months of age, when higher-ranking adult females start harassing them. Such targeted aggression constraints the juveniles’ activities and leads to the gradual emergence of submissive behaviours towards higher-ranking individuals. In turn, the infants of higher-ranking mothers are not targeted in aggressive interactions and as submissive behaviours by subordinates emerge, start initiating conflicts themselves with peers and older individuals. These conflicts are often met with submissive responses due to the threat to the competitor posed by their high-ranking mothers^35^. Therefore, as juveniles age, their probability of winning or losing a conflict is strongly determined by whether their mother outranks their competitor, a process that leads juveniles to obtain rank positions immediately below their mothers. Such behaviourally-mediated rank inheritance has also been demonstrated in natural adoption events of wild spotted hyenas where offspring rank closely matched their adoptive rather than their biological mother^34^. Overall, the different possible pathways of rank inheritance are likely complementary rather than alternative, with prenatal endocrine exposure and genetic inheritance predisposing individuals to be more or less aggressive and postnatal maternal rank shaping rank trajectories. Crucially however, these processes are concerned with offspring living in their natal group and little is known about how maternal rank impacts offspring rank once dispersal takes place.

While in some species it is well-established that maternal rank impacts maternal social behaviour towards offspring, with increased protectiveness associated with lower-ranking mothers^36^, maternal behaviour can still be highly variable even among females of the same rank^37,38^. Individual maternal behaviour may depend on offspring sex^13,39^ as well as on sociecological factors such as the arrival of new group members^40^. In other cases, maternal behaviour directed towards offspring has been found to be remarkably stable over time and contexts. In turn, previous studies across a variety of species have found maternal social behaviour (often referred to as maternal style) to impact offspring social development. For instance, in vervet monkeys, juveniles reared by more rejecting mothers approached and initiated more conflicts with adult males than males reared by less rejecting mothers^38^. In moor macaques (*Macaca maura*), higher maternal grooming was related to a higher probability of immatures engaging in grooming interactions with other partners^41^ and in rhesus macaques, social play initiation by yearlings was negative predicted by maternal cradling^37^.

Yet whether the influence of early maternal behaviour persists into adulthood or is restricted to the juvenile period in the dispersing sex remains an open question. This is due in part to the inherent difficulty of following individuals through dispersal in natural populations. To fill these knowledge gaps, we leverage 16 years of life history and socio-behavioural data to test how maternal rank, maternal centrality, maternal grooming and maternal proximity to sons impact male rank post-dispersal in wild vervet monkeys. Furthermore, given the interdependencies between rank and social behaviour, we test the potential of male grooming and proximity behaviour both pre-dispersal during the juvenile period and post-dispersal, as mediators between maternal characteristics and male rank. We investigate these questions using the long-term data base of the INKAWU Vervet Project (IVP), which includes approximately 600 individually identified wild monkeys from seven habituated groups and a full generation turn-over. Vervets are cercopithecine monkeys with female philopatry and intersexual co-dominance hierarchies^42^ which linearity is ecologically-determined^43^. Maternal rank is inherited by juveniles of both sexes with little maternal intervention in agonistic conflicts^35^ and males disperse upon reaching sexual maturity between four and five years old during the mating season^44^. Consequently, this species represents an ideal system in which to investigate long-term direct and indirect effects of maternal environments on sons rank acquisition because while juveniles inherit their mothers’ ranks in the natal groups and females often maintain them for life, males must integrate into new social environments at least once. Throughout their lifetime, males can disperse multiple times (secondary dispersals) in a process that is relatively understudied and can do so alone as well as in parallel with peers. After dispersal, rank changes are largely socially mediated, with better connection and integration into the group preceding rank increases^45^. Additionally, individuals can also increase in rank via agonistic interactions and after the disappearance of higher-ranking group members but the contribution of these phenomena to the process of rank acquisition remains unknown. The relationship between male rank and mating success in this species is unclear, with some studies in the wild finding no correlation between mating success and dominance rank^46^ while captive studies find that the alpha male sires most of the group’s offspring^47^.

To mitigate the biases intrinsic to field primatology of varying observation efforts across individuals and time periods, we employ a Bayesian framework to build social networks based on grooming and proximity interactions that allows us to translate uncertainty in the observational data into uncertainty around effect estimates in downstream analyses. In addition, we combine this framework with piecewise mediation analyses to investigate potential effects of male social behaviour in mediating the relationship between maternal characteristics and male rank post-dispersal.

While we expected a relationship between maternal and sons’ rank, we had no specific prediction regarding the direction of this relationship. This is because despite the evidence of the silver-spoon effect across multiple species^1^, which would be supported by a positive relationship between sons and mothers’ rank, there is no clear evidence of such effect in the context of rank acquisition^48^. Furthermore, multiple field observations in the target population suggest that sons of higher-ranking mothers attain lower ranks one year after dispersal than sons of lower-ranking mothers (van de Waal pers. comm.). If such observations reflect a general compensatory pattern, we would expect a negative relationship between mothers and son’s rank post-dispersal. Such negative relationship could be explained if upon joining a new group, individuals that had low rank as a juvenile were less likely to initiate or escalate conflicts against established high-ranking residents (which would most likely be lost) than sons of high-ranking mothers, potentially integrating more easily into the group. Conversely, sons of high-ranking mothers might be more likely to challenge high-ranking individuals in the new group, whom they are likely to lose against^49^.

Overall, we expected male social characteristics pre– and post-dispersal to be positively correlated given the consistency in social centrality found in previous vervet studies^50^ and in multiple social behaviours in chimpanzees^51^. We expected this consistency to at least partly mediate any effects found between maternal characteristics and the rank acquired by males in their new groups. However, due to temporal and physical proximity, we expected maternal characteristics to have a stronger effect on males’ juvenile sociality than on post-dispersal sociality. Similarly, we expected males’ social behaviour post-dispersal to have a stronger effect on rank than males’ pre-dispersal sociality. Regarding specific maternal characteristics, we expected sons of higher-ranking mothers to be more social than sons of lower-ranking mothers. Similarly, we predicted that sons that were more groomed and more often found in close proximity to their mothers (irrespective of their rank) would be more social themselves and achieve higher ranks post-dispersal.

By analysing separately maternal rank from maternal investment (grooming and proximity) and centrality, we isolate the effects of maternal social behaviour directed towards sons on males’ rank post-dispersal from social and non-social traits associated with female rank such as access to resources, stress levels or amount of aggression received. Furthermore, by investigating both pre– and post-dispersal social behaviour, we evaluate for the first time the impact that dispersal has on males’ social phenotypes and the carry over effects from juvenile maternal environments on males’ integration and ranks acquired in new social environments.

## Methods

### Ethical considerations

All data were collected following ASAB/ABS guidelines for the care and use of animals. All the behavioural observations of wild vervet monkeys in our study were in accordance with the ethical standards of the University of Lausanne and were approved by the relevant local authority, EzemveloKZN Wildlife, South Africa.

### Study site and population

The data was collected at the INKAWU Vervet Project (IVP), founded in 2010 at the Mawana Game Reserve in the province of KwaZulu-Natal, South Africa (28°00’ S, 31°12’ E). The vervet population monitored by IVP includes approximately 200 individuals in seven neighbouring groups, which are habituated and followed by human observers, with overlapping home ranges. Group sizes ranged between 14 to 81 individuals during the study period (January 2011-April 2025). Individuals in these groups are identified based on unique facial and physical features and visually sexed approximately one month after birth. The vegetation of the area is typical of the savannah biome, consisting of mainly thickets and bushveld. Two distinct seasons occur in a year: a hot, wet season from November to April and a cold, dry season from May to October.

### Data collection and data types

Data was collected by a team comprising both researchers and field assistants (hereafter, observers), who followed the vervet groups eight hours a day up to six days per week. A single group was generally followed by each observer per day with a maximum of three observers per group. To collect data independently, observers had to pass an identification test and an inter-observer reliability test with an experienced researcher (minimum 0.80 Cohen’s kappa^52^). We used four types of data in this study: life-history events, aggressive interactions to calculate rank, grooming interactions, and proximity interactions.

#### 1. Life-history data

Deaths, births, dispersal events and mother identity are regularly recorded in a life-history file. These data were used to identify natal dispersal events of known males between observed groups. We focused on natal dispersals to ensure that all previous social experiences took place in the natal group during the males’ juvenile period.

#### 2. Rank

Conflict data among adults was used to build dominance hierarchies. Conflict data was collected *ad libitum* given the rarity of these events following a predetermined ethogram. For each conflict, the direction and outcome of the conflict were recorded to establish winners and losers. We used only dyadic interactions with a clear ‘winner’ and ‘loser’ to construct the dominance hierarchy. We determined the ‘loser’ of an interaction as the individual that showed a submissive behaviour or a retreat following an aggressive behaviour by another individual (winner). To determine maternal ranks during the juvenile period in the natal group, we extracted the mothers’ ranks based on female-female conflicts from the date of birth of the males until the date of dispersal. We calculated maternal standardized Elo scores using the ‘extract_elo’ function from the ‘EloRating’ package in R^53,54^. We then averaged these scores across the males’ juvenile period. In this way, we generated scores that fall between 0 and 1 allowing comparisons of dominance rank between individuals from groups of varying size. Maternal ranks based on adult female-female conflicts were highly correlated with maternal ranks calculated using conflicts both within and between sexes (p < 0.001, Spearman rho = 0.98).

To calculate the rank attained by each focal male, we computed their rank during their first year after dispersal using male-male conflicts (up to 365 days after their immigration date; median days present first year = 365). If the males left the group before the one-year mark, they were not excluded from the analyses (as this could have biased the results). In these cases, we used all their available data post-dispersal. In turn, we included in all models the number of days males stayed in the immigration group during the first-year post-dispersal to account for differences between males in observation effort. Specifically, this value was included as a predictor of the residual standard deviation (σ), meaning that the precision of the fitted relationship between post-dispersal social network metrics, maternal characteristics and male rank varies according to how long males stayed in their group post-dispersal. The one year time point was chosen based on previous studies which found that this is the median time period needed to reach maximum rank in other populations of this species^44^. This relatively short timeframe also allowed us to focus on the integration phase post-dispersal. Males included in the rank analyses had between 2 and 184 male-male conflicts^44,55^ recorded in the first-year post-dispersal (median conflicts = 14.5). Excluding males based on an arbitrary number of conflicts could have biased our sample towards males that either initiated or where targeted more often in aggressive interactions in their new groups during their first-year post-dispersal, omitting data from males that might have been adept at avoiding conflicts. We thus decided to include data from all males from which a sequence of conflicts could be established based on at least 2 conflicts. Male ranks one-year post-dispersal based on both within– and between-sex conflicts were highly correlated with ranks calculated from male-male conflicts (p < 0.001, Spearman rho = 0.78). We used a year of conflicts from the groups males immigrated into prior to each males’ immigration date to allow for a stable hierarchy to form into which dispersing males integrated.

#### 3. Grooming

We used grooming data collected *ad libitum*^56^ to build males’ affiliative networks pre– and post-dispersal. In each grooming interaction, the identity of the individuals involved in the behaviour as well as the direction of the behaviour were recorded. A grooming bout starts when one individual start grooming another and ends when the groomer stops for more than one minute, moves at least two meters from the groomee, the groomee takes over, or the groomer changes partner. Event duration is not recorded to allow observers to record as many group members simultaneously as possible. Observers are trained to reduce sampling bias by moving throughout the group during data collection rather than standing at a single observation point. While grooming interactions can also be collected via other observational methods such as focal sampling, which allow behavioural metrics to be standardized by observation effort, focal sampling was only implemented at IVP in 2022, meaning that the volume of data available is limited. Furthermore, previous studies using IVP data found that behavioural networks built using *ad libitum* and instantaneous focal sampling every two minutes during 20 minute focal follows were equivalent^57^. The IVP data collection protocol for grooming interactions follows established protocols in other long-term field sites^58^ which aim is to maximize the number of events recorded. We further accounted for differences in sampling effort through time and across individuals by building Bayesian directional grooming networks where edge weights correspond to the proportion of observation days a dyad was observed grooming (see below). Pre-dispersal metrics were calculated using data from the first birthday of males (when they are almost fully independent from their mothers) until their dispersal. Post-dispersal metrics were calculated from the time the males were seen for the first time in their new group until one-year post-dispersal at the latest.

For each focal male, we also calculated the amount of grooming received from its mother during their first year of life as a measure of early maternal investment. This metric was considered as independent from maternal rank as they were uncorrelated both in this data set (r = 0.04, p = 0.76, n = 70) and previous studies in the same population^13^. Grooming received was computed as the proportion of grooming events that a mother directed to her son out of all grooming events the mother was involved in as a groomer. Focal males were observed in grooming interactions a median of 191.5 days post-dispersal (range 21 – 294), a median of 695 days pre-dispersal (range 194 – 1156) and were observed being groomed by their mothers a median of 41 days (range 0 to 284).

In addition, we calculated the eigenvector centrality in grooming networks of mothers during the juvenile periods for the focal males. As expected, provided that high-ranking individuals generally occupy central positions in their groups’ social structures, this metric was positively but weakly correlated to the females’ rank (r = 0.24, p = 0.04, n = 70).

#### 4. Proximity data

We used proximity data collected via scan sampling to build proximity networks. Before focal follows were implemented at IVP in 2022, scans were conducted every 30 minutes for up to 10 minutes. Since 2022, two scans of up to 30 minutes (morning and afternoon) were conducted starting at the full hour. This protocol was inspired by the data collection protocols of the long-term Amboseli Baboon Project and the Cayo Santiago Project where proximity data is collected sequentially as a snapshot for each individual in the group^58^. While in Amboseli such scans are open ended, IVP scans (like the scans in Cayo Santiago) are limited in time to ensure that the context in which the proximity data is collected is equivalent for all individuals sampled. If the behavioural context changed during a scan (due to e.g., an alarm call, a between-group encounter, a river or road crossing), the scan was terminated. During each scan, observers recorded the identity of all individuals within 1m (until 2022) and within arm length (from 2022 following a protocol change) for as many individuals in the group as possible. Both distances were pooled given that they are both non-contact, proximity distances differing in less than 50 cm. When comparing the individual metrics calculated from networks built from each of the two distances (1m and arm length) on those years where both distances were collected (2016-2021), we found a strong correlation in the number of proximity partners (Spearman rho 0.83, p < 0.001) and the strength of proximity connections (Spearman rho 0.69, p < 0.001). For each dispersing male during their juvenile period and their first-year post-dispersal, we built separate Bayesian weighted undirected proximity networks (see below). Males were sampled in a median of 102.5 proximity scans (range 1 to 471) post-dispersal and in a median of 571 scans (range 104 – 912) in their natal group from their first birthday until the time they dispersed.

As a second metric of maternal investment complementary to grooming directed towards sons, we calculated the proportion of scans during the first year of a males’ life in which they were observed within five meters of their mothers. This metric was also considered as independent from maternal rank as they were uncorrelated (r = –0.21, p = 0.09, n = 70). Proximity to sons and grooming of sons were significantly and negatively correlated, albeit with a low correlation coefficient, and thus were used as complementary predictors potentially representing a trade-off in maternal investment behaviours (r = –0.26, p = 0.019, n = 78). Only the first year of life was considered as infant identity is not recorded during scans (only their presence) and after a year individuals are considered juveniles rather than infants and mothers could have other dependent offspring. Focal males were observed in a median of 197 scans within five meters of their mother their first year of life (range 4 to 575).

We also calculated maternal eigenvector centrality in proximity networks using the same approach as described for the maternal eigenvector centrality in grooming networks above. As expected given the high centrality of high-ranking individuals within groups, females’ eigenvector centrality in proximity networks was significantly but weakly correlated with their rank (r = 0.38, p < 0.001, n = 70).

### Statistical analyses

To investigate direct and indirect effects of maternal characteristics on male rank and the role of social behaviour in mediating this relationship we combined Bayesian social network estimation with mediation analyses.

### Network estimation

For each immigrant male, we constructed proximity and grooming Bayesian networks both pre-dispersal and one year post-dispersal. To build the undirected proximity networks, we first generated a posterior distribution of dyadic indices using a Bayesian conjugate approach with Beta(0.1+ *x*, 0.1+ *y*) priors, where *x* is the number of co-occurrence scans and *y* is the number of scans in which either or both individuals were observed (equivalent to the Simple Ratio Index^reviewed^ ^by^ ^59^). From each of these posteriors, we then built edge weight distributions by drawing 1000 samples. To build directed grooming networks, we first generated a posterior distribution of the proportion of days a dyad was observed grooming (*x*) out of the total number of *ad libitum* observation days conducted in the group (*y*) using the same Beta(0.1, 0.1) priors as before. Edge weight distributions were then built by drawing 1000 samples from these posteriors. An equivalent approach was used to build the maternal networks from which we calculated maternal centrality in the grooming and proximity networks.

For each male and for each of the 1000 posterior draws we calculated the following network metrics pre and post-dispersal: a) proximity degree: number of individuals the male was observed in proximity with; b) proximity strength: sum of all edge weights in the proximity network; c) grooming in-degree: number of individuals the male was observed being groomed by; d) grooming out-degree: number of individuals the male was observed grooming; e) grooming in-strength: sum of all incoming edge weights in the grooming network; f) grooming out-strength: sum of all outgoing edge weighs in the grooming network. The pre-dispersal degree metrics were normalized by the total number of individuals present in the groups across the multi-year juvenile period minus one whereas the post-dispersal degree metrics were normalized by the median daily group size minus one. This difference was due to the variability in group size across years for the pre-dispersal period which, contrary to the post-dispersal period, included multiple birth and migration seasons.

### Mediation Analyses

We used mediation analysis (Fig. 1) to investigate the potential role of male social behaviour pre– and post-dispersal as a mediator of the relationship between maternal characteristics (rank, grooming of sons, proximity to sons and network centrality) and male rank in the first-year post-dispersal. The six social network metrics described above (proximity strength and degree; grooming in– and out-strength and degree) were each tested separately as mediators. For each maternal characteristic–mediator combination, we fit a joint Bayesian multivariate regression model, estimating simultaneously: (a) the path from the maternal characteristic to mediator; (b) the path from the mediator to male rank, controlling for the maternal characteristic; and (c’) the direct path from the maternal characteristic to male rank, controlling for the mediator. Because paths a and b were estimated within a single joint model, their posterior draws were paired, allowing the indirect effect (a × b) to be computed as a posterior distribution reflecting the joint uncertainty in both paths. The total effect (c = c’ + a×b) was additionally estimated as a simple regression of male rank on the maternal characteristic alone, with no mediator included. Each model was fit separately to 1,000 posterior draws of the mediator (and, where the predictor was itself network-derived, of the predictor), drawn jointly from the relevant network posterior to propagate network uncertainty. Posterior samples from all 1,000 draws were pooled into a single combined posterior distribution per path, from which we report the posterior mean and 95% credible interval (2.5th–97.5th percentile). All variables were standardised (z-scored) prior to analysis.

**Figure 1:**
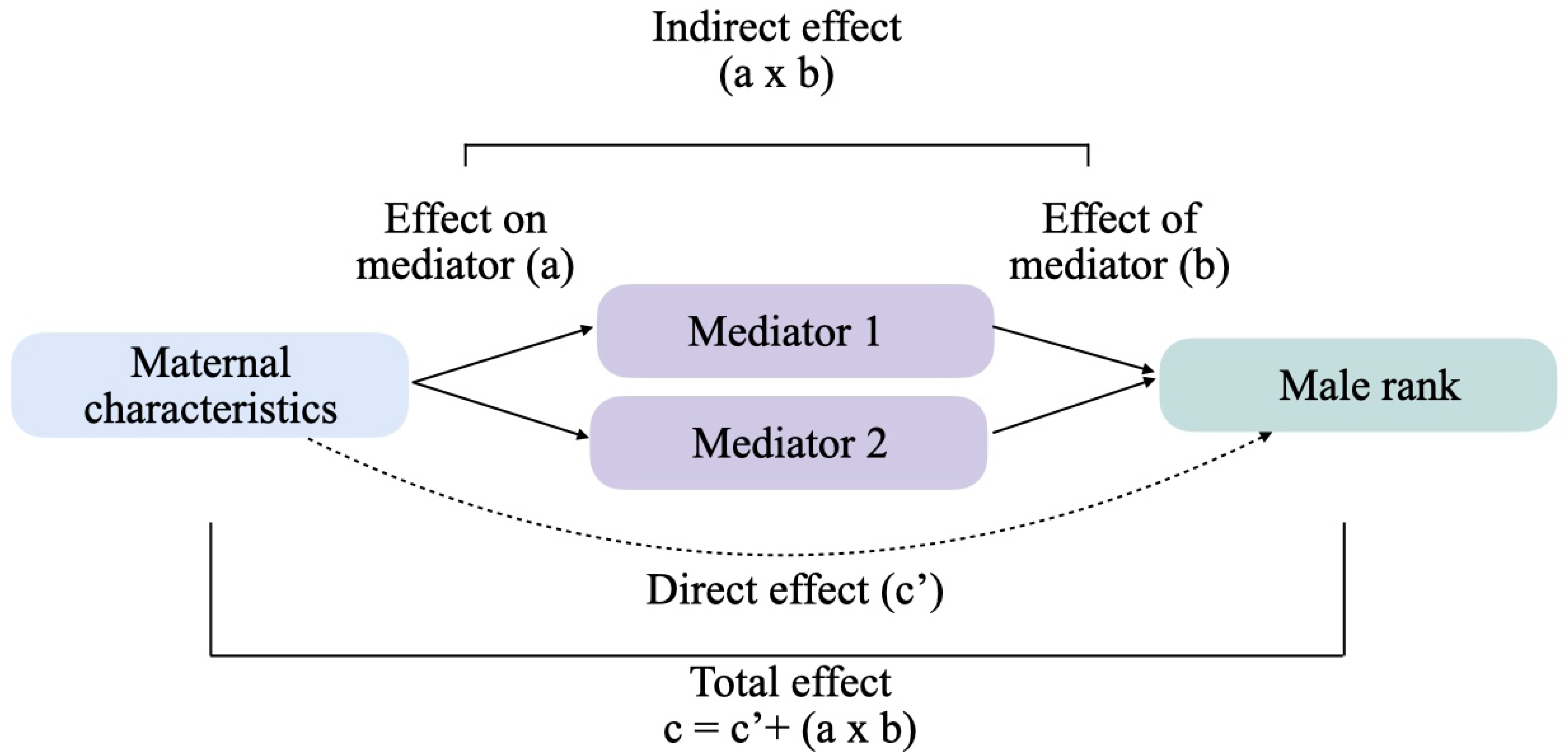
Schematic representation of the mediation analysis. The total effect (c) of maternal characteristics on male dominance rank acquired one-year post-dispersal was decomposed into a direct effect (c’) and indirect effects operating through social network metrics (mediators). Path a represents the effect of the maternal characteristic on each mediator. Path b represents the effect of each mediator on male rank, controlling for the maternal characteristic. The indirect effect (a × b) quantifies the portion of the total effect transmitted through each mediator. Two versions of the analysis were conducted: one using as mediators social network metrics from the pre-dispersal period and one using as mediators social network metrics from the first-year post-dispersal.

### Group characteristics

We conducted an additional analysis to evaluate how much variation in the maternal characteristics and juvenile social metrics was explained by the identity of the natal group, given the known differences in sociality among groups in the study population^60^. We ran a Bayesian random-intercept model where we first partitioned variance in maternal characteristics and sons’ juvenile network metrics into within– and between-natal-group components. Then, we derived the intraclass correlation (ICC) as the proportion of variance attributable to natal group identity, refitting across 1,000 posterior draws per male where the outcome was itself a posterior-derived network metric.

We also quantified how much of the variation in the males’ social network metrics post-dispersal and rank was explained by the demography of the immigration group. To do so we ran two Bayesian linear regression models testing whether destination-group size and adult sex ratio at the time of immigration predicted males’ characteristics post-dispersal.

## Results

We found 212 males which were born in the population and performed their natal dispersal among groups habituated to human observation. Of these, we were able to calculate maternal ranks during the juvenile period, males’ rank during the year post-dispersal and both males’ proximity and grooming network metrics post-dispersal from 33 males. The analyses testing maternal investment effects and maternal centrality on male rank mediated by males’ sociality post-dispersal included 38 males. The analyses testing males’ juvenile social behaviour as mediator between maternal characteristics and males’ rank post-dispersal included 36 males. The reduction in the number of males available in our sample was mostly due to males having unknown mothers, no recorded date of birth, the mothers not having any recorded conflicts in the natal group which prevented the calculation of maternal ranks or males not staying long enough in their immigration groups in order to have recorded conflicts. Males dispersed when they were a median of five years and one month (range: four years and five months to six years and one month).

### Maternal effects on juvenile sociality

Multiple maternal characteristics were found to influence males’ juvenile social behaviour (a-paths). Maternal grooming investment during the first year of life was positively associated with all metrics of sons’ sociality. Sons’ grooming in-strength (β = 0.474, 95% CI [0.158, 0.789], probability of direction range = 99.8%), out-strength (β = 0.405, 95% CI [0.078, 0.732], pd = 99.2%), in-degree (β = 0.437, 95% CI [0.119, 0.754], pd = 99%) and out-degree (β = 0.502, 95% CI [0.194, 0.810], pd = 99%) were all positively related to the amount of maternal grooming they received. This indicates that infants of more grooming mothers were more integrated in the grooming network and more tolerated by group members as juveniles. Importantly, these grooming effects were not driven by the grooming relationships between the sons and their mothers directly during the juvenile period as excluding the mother-son interactions did not alter the son’s grooming metrics meaningfully (e.g., in-strength with and without maternal interactions: r = 0.99; out-strength: r = 1). Similarly, males’ proximity degree and strength were also positively related to maternal grooming investment (strength: β = 0.505, 95% CI [0.181-0.830], pd = 99%; degree: β = 0.520, 95% CI [0.226, 0.817], pd = 99%).

Contrary to maternal grooming, the frequency of maternal proximity to sons was negatively associated with all juvenile grooming metrics (in-strength: β = –0.467, 95% CI [-0.778, –0.153], pd = 99.8%; in-degree: β = –0.501, 95% CI [-0.801, –0.198], pd = 99.9% out-degree: β = –0.603, 95% CI [-0.886, –0.306], pd = 100%; out-strength: β = –0.514, 95% CI [-0.827, –0.198], pd = 99.9%). This suggests that higher maternal proximity to sons reduced their engagement in grooming interactions and their integration in the grooming network. Maternal proximity did not impact males’ proximity metrics (Fig. 2).

**Figure 2.**
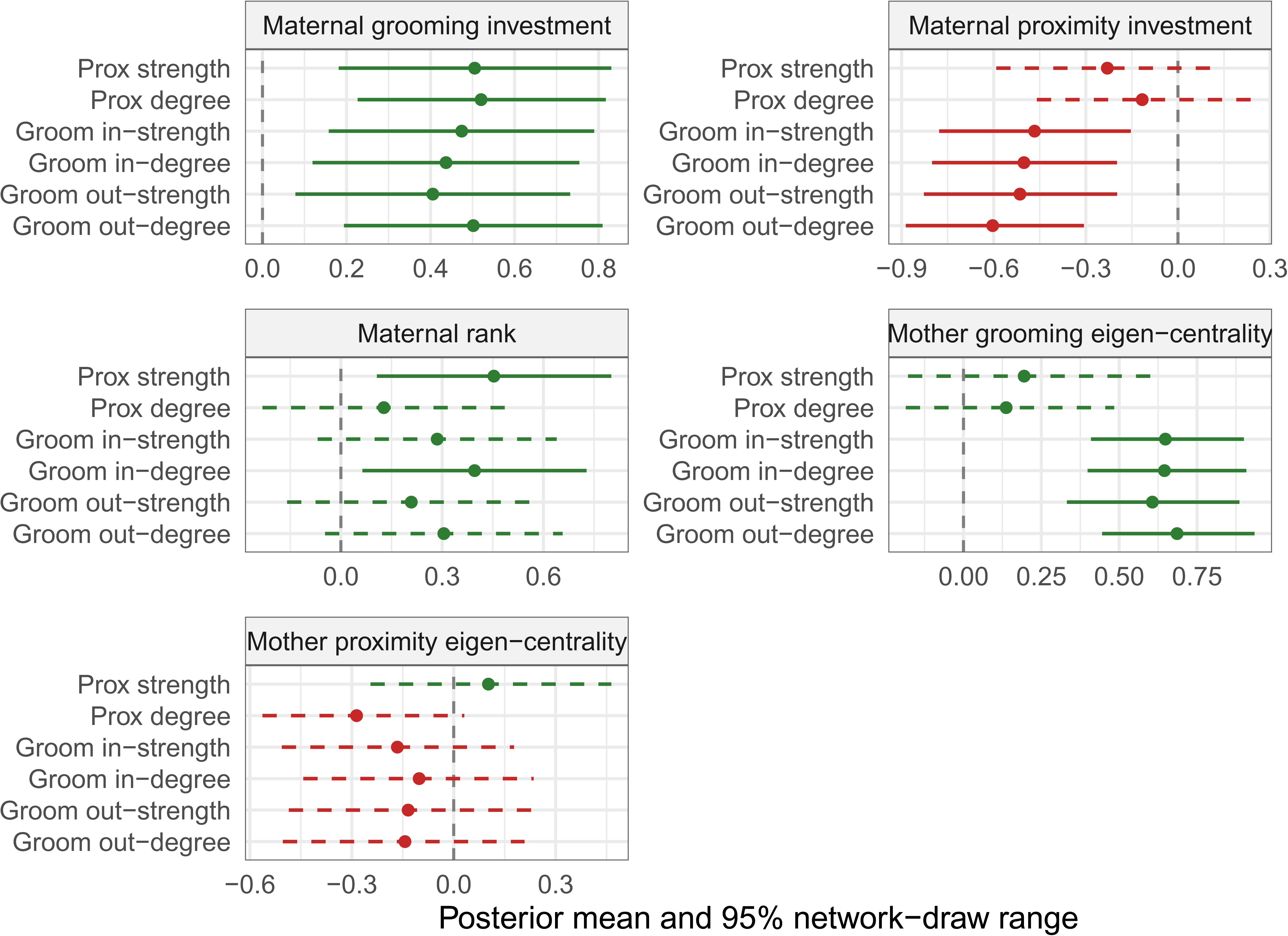
Posterior mean associations between maternal characteristics and sons’ social network metrics during the juvenile period (a-paths). Points show the posterior mean; bars show the 95% network-draw range (2.5th–97.5th percentile across network reconstructions). Solid lines indicate associations for which this interval excludes zero; dashed lines indicate associations for which the interval includes zero. Colour denotes the direction of the point estimate (green = positive, red = negative).

Maternal rank was positively associated with sons’ juvenile proximity strength and with the amount of grooming sons received, with sons of higher ranking mothers being found more often in close proximity to conspecifics and being more groomed by other groom members (strength; β = 0.453, 95% CI [0.107, 0.801], pd = 99.4%, n = 33; in-degree: β = 0.396, 95% CI [0.064, 0.728], pd = 98.9%). The mothers’ own grooming centrality had positive effects on all grooming metrics: in-degree (β = 0.645, 95% CI [0.399, 0.908], pd = 99%), in-strength (β = 0.648, 95% CI [0.409, 0.900], pd = 99%), out-degree (β = 0.686, 95% CI [0.445, 0.935], pd = 99%), and out-strength (β = 0.607, 95% CI [0.332, 0.886], pd = 99%). Mother’s centrality in proximity networks was not related to any of their sons’ social network metrics pre-dispersal (Fig. 2).

None of the 95% credible intervals of the b-paths (effects of social network metrics during the juvenile period on male rank post dispersal, Fig. 3) nor the direct effects (c′ paths) of maternal effects on male rank post-dispersal excluded 0 in any pre-dispersal model.

**Figure 3:**
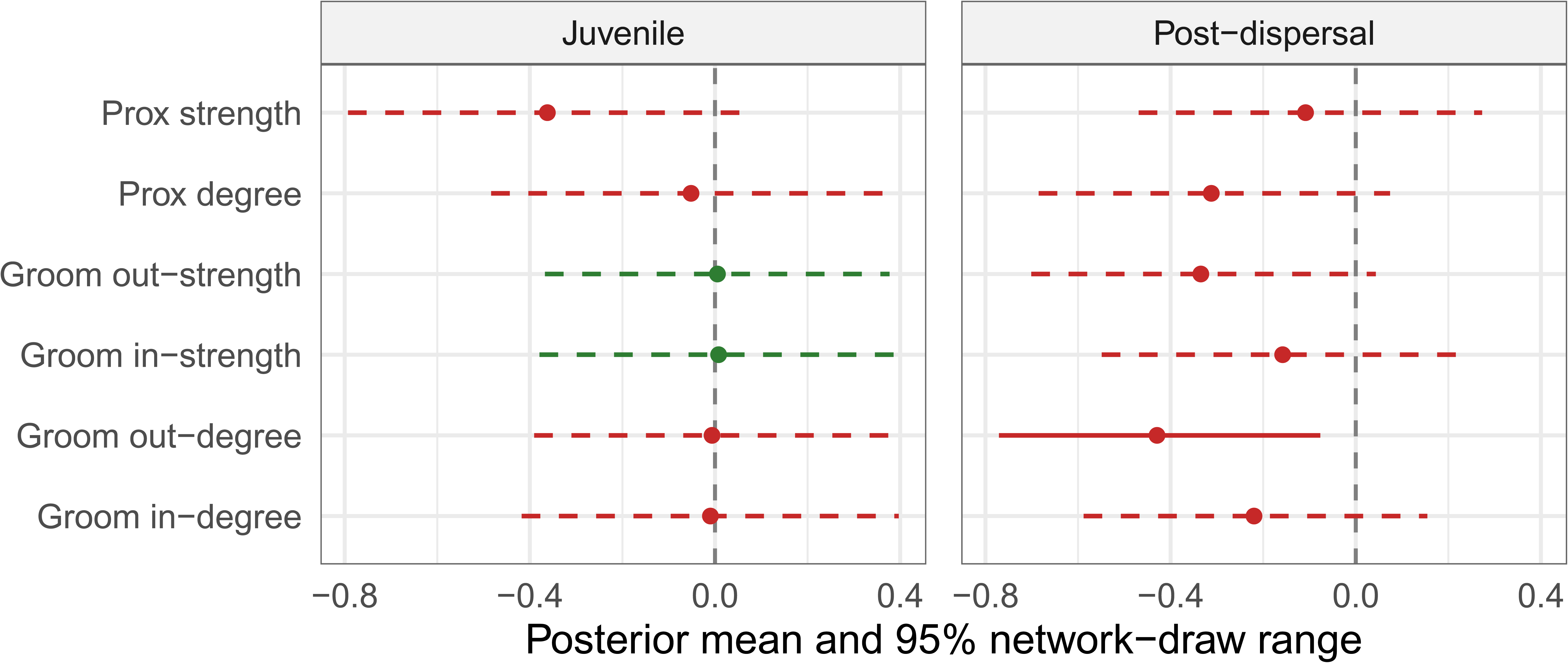
Effects of juvenile (left) and post-dispersal (right) social network metrics on male rank post-dispersal (b-paths). Points show the posterior mean; bars show the 95% network-draw range (2.5th–97.5th percentile across network reconstructions). Solid lines indicate associations for which this interval excludes zero; dashed lines indicate associations for which the interval includes zero. Colour denotes the direction of the point estimate (green = positive, red = negative). N = 33 males in both analyses.

### Effects of post-dispersal social behaviour on male rank

Across models (Tables S1-S5), we found a consistently negative b-path from males’ social characteristics post-dispersal to male rank. However, we only found a single path where the 95% credible interval of an effect excluded 0 (Fig. 3). This was in the maternal rank model, where male’s grooming out-degree was negatively related to males’ rank (β range = –0.43, 95% CI range [-0.77, –0.08], pd = 99%). This result indicates that once males disperse, those males that groom a higher number of partners have lower ranking than the males that groom a lower number of individuals.

Among the a-paths quantifying the relationships between maternal characteristics and males’ social behaviour post-dispersal, the strongest directional signals — though neither excluded 0 from the credible interval — was maternal rank’s negative association with son’s post-dispersal proximity degree (β = −0.318, 95% CI [−0.67, 0.03], pd = 96.3%). Similarly, there was a trend for a mediation effect of proximity degree between maternal and male rank (β = 0.099, 95% CI [−0.03, 0.31], pd = 91.6%). None of the 95% credible intervals of the direct effects (c′) between maternal and male rank excluded 0 in any model (Tables S1-S5). Yet, the four males that achieved alpha positions in their immigration group’s dominance hierarchies were sons of low (n = 1) or middle-ranking mothers (n = 3).

### Relationship between pre– and post-dispersal sociality

Grooming network metrics showed consistently positive (Fig. 4), albeit moderate, correlations between the juvenile pre-dispersal and post-dispersal periods: in-strength (r = 0.486, n = 46), out-strength (r = 0.374), in-degree (r = 0.188), and out-degree (r = 0.152). Proximity metrics were less consistent pre– and post-dispersal. The range of correlation values for proximity strength included 0 (Fig. 4) and proximity degree was the only metric negatively correlated between the juvenile and post-dispersal periods (r = –0.212, n = 46).

**Figure 4:**
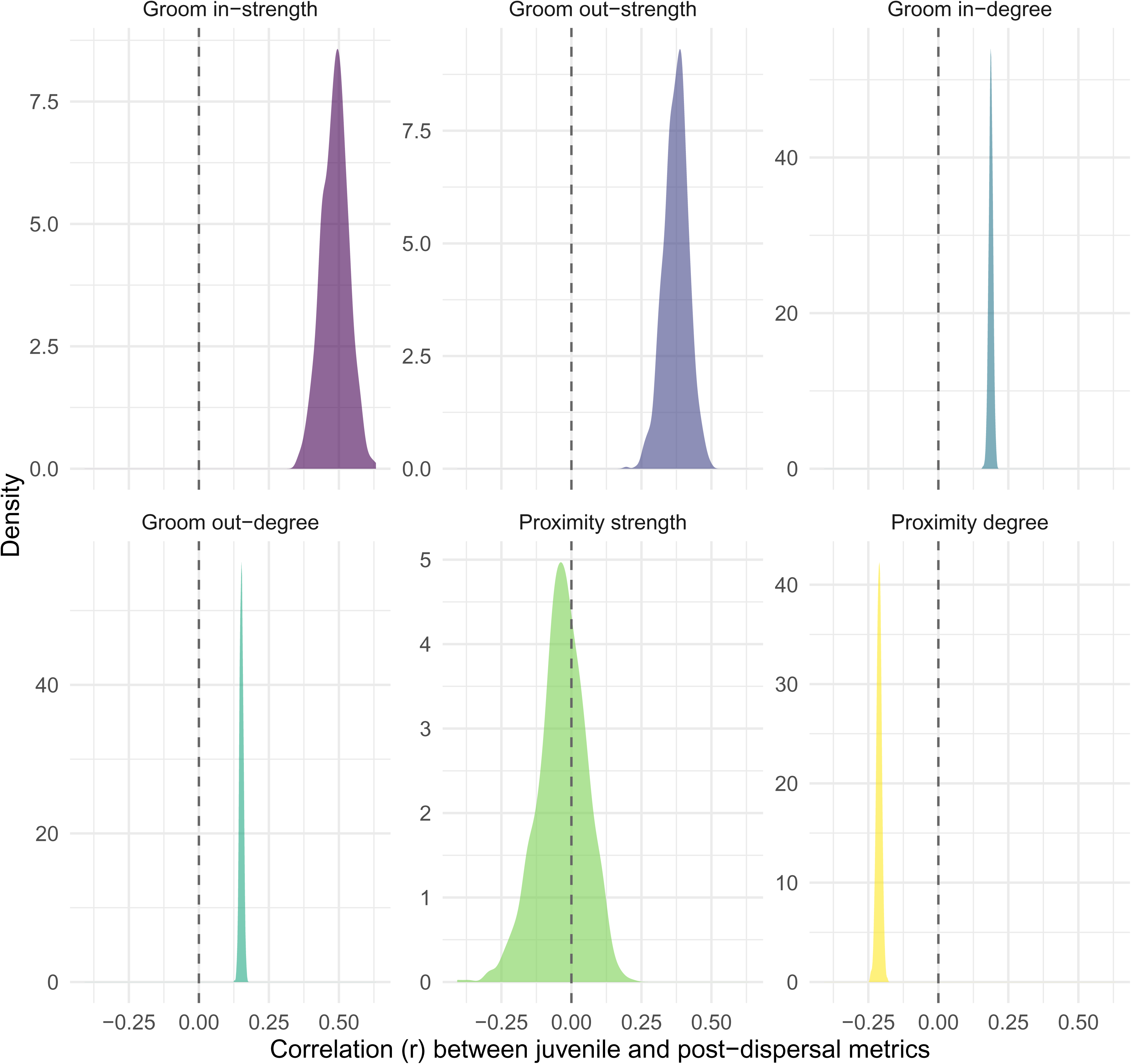
Distributions of the posterior correlation (Pearson’s *r*) between males’ juvenile-period and post-dispersal social network metrics, propagated across 1,000 alternative Bayesian network reconstructions (n = 46).

### Natal and Immigration group effects on predictor and mediator variables

Natal group identity explained very little variance in maternal characteristics (posterior median ICCs range 0.05-0.10). For juvenile social network metrics, natal group identity explained part of the variation in proximity metrics among males (median ICC strength = 0.72, 95% CI [0.26, 0.97]; median ICC degree = 0.66, 95% CI [0.23, 0.96]), but it is important to consider that the different groups were unevenly represented in the conservative subsample of 33 individuals used for this test (range of 3 males from LT and 15 from AK, Figure 5). Males born in KB (n = 4) showed the highest strength of proximity relationships and males born in NH the lowest, whereas the number of proximity partners was highest in AK and KB (posterior mean ∼0.58). The four juvenile grooming metrics were not credibly influenced by natal group identity (median ICCs 0.07-0.31).

**Figure 5:**
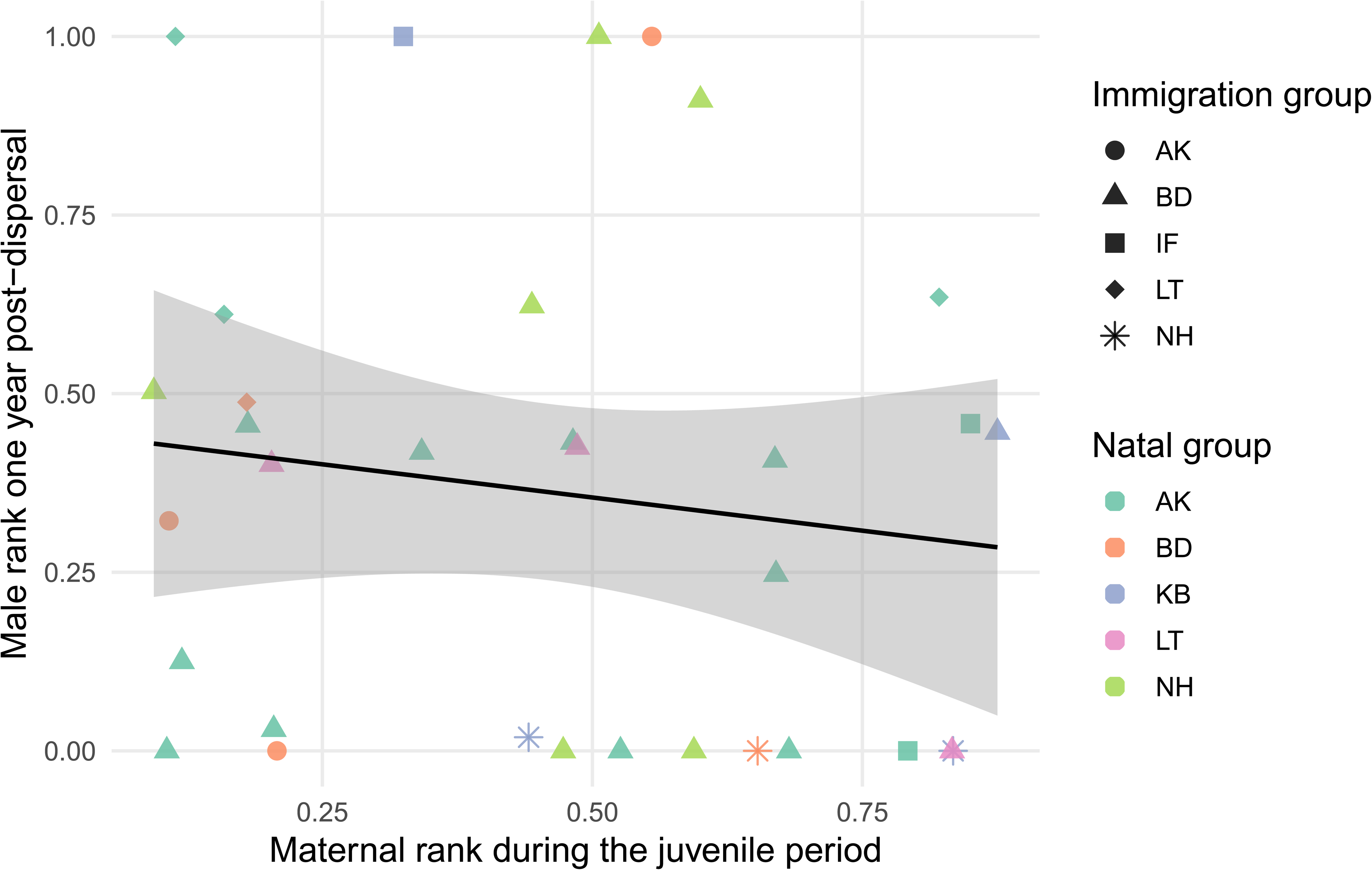
Relationship between maternal and male rank calculated via elo-ratings of all 33 males included in the maternal rank mediation analyses. Point shapes represent the different groups males immigrated into whereas point colours represent the males’ natal groups. The black line represents the linear model with the grey ribbon representing the 95% confidence interval around the estimate.

The size of the group’s individuals dispersed into at the time of immigration did not show any significant effect on males’ rank or social network metrics post-dispersal. However, adult sex ratio at the time of immigration was positively associated with all four grooming-based network metrics (regression coefficients’ range 0.34-0.56 or change in metric for each additional female), but not the proximity metrics. This means that males who immigrated into groups with a higher female to male ratio had higher numbers of grooming interactions and partners during their first-year post-dispersal. However, this could be explained by many of the males in the sample immigrating into the BD group (Fig. 5), which is the largest in the population and has a high female-biased ratio.

## Discussion

Using 16 years of demographic, life-history, and behavioural data from wild vervet monkeys, we find that while males’ social behaviour post-dispersal is partly related to their rank, contrary to our predictions maternal characteristics during the males’ juvenile period do not directly affect males’ behaviour nor rank once they disperse. Instead, maternal characteristics are associated with males’ juvenile social behaviour, which is only moderately consistent pre– and post-dispersal.

Previous studies in the same population found that sons of higher ranking mothers had more partners and stronger relationships in the natal group than sons of lower ranking mothers^61^. In the present study we partly replicated these results, finding that the number of proximity associations of males during the juvenile period was positively related to maternal rank. This effect could be due to higher ranking mothers facilitating the integration of males by exposing them to their own social environments. However, the lack of effect of maternal proximity and grooming centralities on juveniles’ proximity strength suggests that maternal rank is related to sons’ social proximity to other group members via other mechanisms beyond social facilitation (see also^13^). Rank-modulated aggression represents an alternative mechanism explaining his relationship. Sons of high-ranking mothers experience less aggression and displacement from conspecifics^35^. In turn, this lower degree of harassment might also be related to sons of high-ranking mothers generally having better access to resources and thus being physically larger^62^. Combined, these factors could lead sons of higher-ranking mothers to be tolerated in close proximity more often by conspecifics, as well as potentially increasing the boldness of these higher-ranking sons.

Also in agreement with previous studies in the same population^13^, we found that maternal rank was positively related to juvenile grooming in-degree, meaning that sons of higher ranking mothers were the recipients of grooming by a broader set of group members than sons of lower ranking mothers. However, we did not find the broader effects on grooming reported by Tanking *et al*^13^ who compared maternal effects on male and female offspring juvenile fitness and social behaviour. The differences between studies could be due methodological differences as the analytical approach used, the social network metrics calculated, and the samples of males vary between the two studies (400 versus approximately 40 in the present study). Previous studies in rhesus macaques, similarly to our results, found no effect of maternal rank on juvenile’s own grooming engagement^63^.

We found opposing effects of different maternal behaviours directed towards sons during their first year of life on their sociality as juveniles. Individuals that received more maternal grooming provided and received more grooming interactions themselves. Such results are similar to those found in Moore macaques^41^, where sons of mothers that groomed their sons more often engaged in more social behaviours themselves like social play. Furthermore, we found that sons of more central mothers in the grooming network had higher grooming engagement with more individuals as juveniles. This overall association between maternal and juvenile grooming could indicate that male vervet monkeys learn or inherit grooming tendencies from their mothers via social inheritance or social scaffolding. Social learning, both of interactions directed towards infants as well as observed, is likely to contribute to the development of social behaviour in juveniles given its role in the acquisition of social conventions in primates^64^. In addition, maternal grooming could also shape offspring physiology which can in turn impact individuals’ social behaviour. Receiving more maternal grooming has been found to reduce stress levels and confer higher stress resilience in primates^65^ and rodents^66^ which might facilitate the integration of the juveniles in their social group. Such hypotheses are supported by laboratory studies where individuals pharmacologically treated with anxiolytics engaged in more social behaviours including grooming and close proximity to conspecifics^67^.

Contrary to our grooming results, maternal proximity to sons during the first year of life was negatively related to the number of grooming partners juveniles had as well as the amount of grooming juveniles provided. These results suggest that while maternal grooming fosters integration into the grooming network, frequent proximity to the mother (without grooming) might restrict juveniles to their mothers’ immediate social circle, preventing sons from developing social relationships with non-kin or individuals not closely affiliated to their mothers. This latter result could be due to mothers physically restricting or limiting their sons’ social interactions as proximity and restraint are generally clustered as a part of a protective maternal style^36,38^, but we did not record this type of interactions and thus this hypothesis remains to be tested. The centrality of mothers in their group proximity networks was generally negatively related to juvenile males’ grooming behaviour but the posterior distributions of these effects included 0 in all cases. These results contrast those found in other species with male-philopatry such bonobos, where it has been proposed that mothers help sons occupy desirable spatial positions within the group that allow males to closely interact with females and thus, positively affect their mating opportunities^68^.

Previous studies have suggested that grooming and proximity interactions fulfil different social roles^69^. Proximity relationships have been argued to reflect shared use of resources, information sharing and social tolerance^70^, whereas grooming interactions represent behavioural commodities^71^ that serve as the main social currency in primates and affect multiple aspects of individual’s physiology^72^ and brain activity^73^. The moderate correlation or absence altogether between metrics calculated from both types of relationships in previous studies further supports this distinction^74,75^. Our findings that maternal proximity and grooming differently affect males’ juvenile proximity and grooming behaviour further support this conclusion. These functional differences between social behaviours are also reflected in our post-dispersal results.

We find that grooming more individuals, but not proximity, is associated with lower ranks after first dispersal. This result is consistent with previously reported subordinate-to-dominant grooming patterns in primates^76,77^, where grooming is traded for rank-related benefits such as feeding tolerance^15^. This result is also likely to reflect differences in selectivity among males of different rank, with lower ranking individuals grooming more opportunistically and higher-ranking individuals being more selective in their grooming partners (e.g., higher ranking or central females from the dominant matriline). Broadly, the fact that lower ranking individuals engaged with more grooming partners empirically validates previous theoretical predictions suggesting that individuals prioritize investing in quantity-based social strategies to increase their access to monopolizable resources^78^.

Our null total effect of maternal rank on male rank post dispersal is in agreement with previous vervet studies in captivity. Fairbanks et al.^48^ found maternal and sons’ vervet adult rank to be uncorrelated after males were transferred to new groups simulating dispersal events with impulsivity and better body condition leading to higher ranks. Our study now extends this finding to wild populations. The same pattern that we find in vervets has also been found for mandrills^79^ and baboons^80^, where maternal rank influences were only relevant during the juvenile but not the adolescent or adult stages. However, we also made the interesting observation that all four males that attained alpha positions in the groups they immigrated into during the first year if life, were sons of low or middle-ranking mothers which corroborates our field observations. Future research should explore this potential pattern further with a larger data set. In addition, future research should assess whether rather than maternal characteristics, male rank post-dispersal is more strongly determined by personality traits, male services provided in the new group such as alarm calling^81^ or between group conflict participation ^82^, bonds with influential females ^21^, or body condition at the time of dispersal^48^.

Our results on the correlations among social behaviour across life-stages — moderate consistency in grooming behaviour but not in proximity — partly parallel previous findings that vervet monkeys are consistent in grooming-based network measures across seasons within a stable social group^83^. Our results extend this pattern to not only multiple seasons but a complete change in social environment, group membership, and available social partners following natal dispersal. We find that grooming-based social tendencies persisted across this transition, while proximity associations did not. This difference could be due to active social behaviours (grooming) being more stable individual traits than spatial associations (proximity). Proximity relationships, as a passive interaction, are likely to be shaped by the specific social opportunities available in a male’s new group rather than a persistent individual disposition. Previous studies in female baboons have demonstrated that adult grooming tendencies are moderately heritable^84^, which could explain both the relationship between maternal grooming and male juvenile grooming and the moderate correlation between pre– and post-dispersal grooming. However, the fact that we do not find that maternal behaviour during the juvenile period extends to the post-dispersal period directly suggests that this influence might be too subtle to be detected with the available sample size from our study population.

Finally, previous studies on the target population revealed group differences in sociality that could have influenced our results both in terms of the males’ natal groups and their immigration group^60^. Natal group identity explained little of the variance in maternal characteristics and juvenile grooming but there were differences in the frequency of proximity relationships and the number of proximity partners among males born in different groups. Regarding the groups males immigrated into, 54% of males dispersed into the largest group in the population (BD), meaning that the immigration context was relatively similar across males. Thus, we could not explore the effect of group identity directly on post-dispersal metrics given the low diversity of target groups. Instead, we explored how the specific characteristics of the groups at the time of immigration impacted males’ rank and social behaviour. Group size did not explain almost any of the variation in the target metrics whereas sex ratios had a larger effect in some cases. Specifically, sex ratio was positively related with grooming, with all grooming metrics having larger values the larger the number of females relative to males in the immigration groups. However, given that many of the highest sex ratios were found at different points in time in the BD groups, which is the largest in the population, we cannot exclude that this association is group specific.

Overall, our study highlights the value of combining large scale data sets generated by long-term field sites with Bayesian network approaches and mediation analyses to unravel the different pathways and timescales at which early maternal environments shape the social behaviour of the often-overlooked dispersing sex. We provide the first evidence of the absence of a link between maternal and male rank post-dispersal in wild vervet monkeys. We also reveal that while maternal environments are related to males’ social behaviour during the juvenile period, these effects do not extend to males’ social behaviour post-dispersal despite moderate within-individual correlations in grooming predispositions. Taken together these findings provide new insights into the extents and limitations of early life maternal characteristics on male social behaviour before and after the crucial life transition in primates that is the integration into a new social group.

## Acknowledgments

We thank the IVP onsite managers, Albert Driescher, Arend van Blerk, Michael Henshall, Nokubonga Dlamini, Siboniso Thela, Zonke Mbutho and all the field assistants, Master students, PhD students and postdocs who collected the data over the study period. We are grateful to the van der Walt family for giving us the permission to conduct the study on their land. This project was funded by the Swiss National Science Foundation (P300P3_151187, 31003A_159587, PP00P3_170624, PP00P3_198913 and CRSII-222818) along with Branco Weiss Fellowship–Society in Science, the grant ‘ProFemmes’ of the Faculty of Biology and Medicine, University of Lausanne and by the European Research Council under the European Union’s Horizon 2020 research and innovation programme for the ERC ‘KNOWLEDGE MOVES’ starting grant (grant agreement no. 949379) that also supported E.v.d.W. during the time of writing. We are grateful to Josefien Tankink and two anonymous reviewers for feedback on previous versions of the manuscript.

## Code availability

All code used to analyze the data included in this manuscript is available at https://github.com/AlbaMotes/vervet_integration_postdispersal.git. The raw data files used in the present manuscript are deposited in the Zenodo repository 10.5281/zenodo.23532870.

## Supplementary Material

**Table S1:**
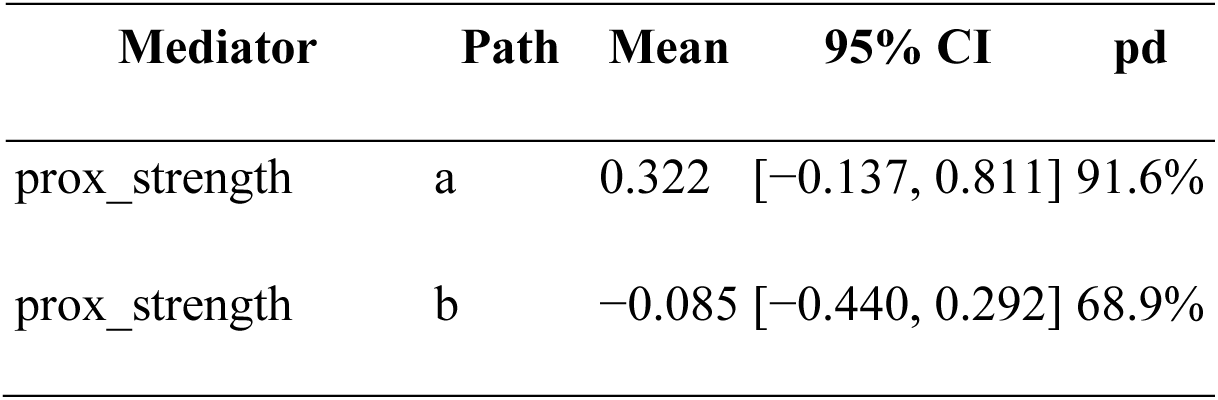
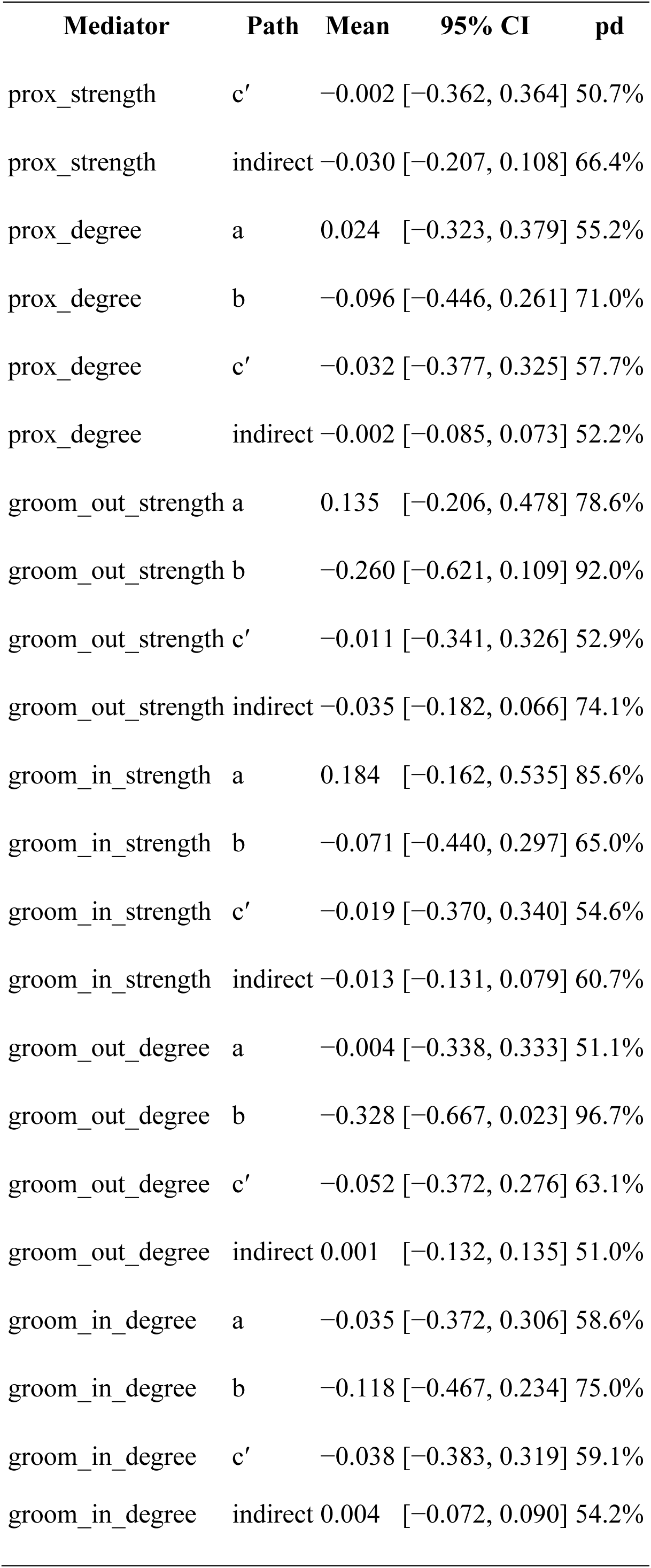
Posterior mean, 95% credible interval (equal-tailed quantile), and posterior probability of direction (pd = the larger of prop_positive/prop_negative) for every path in the model testing the effect of maternal grooming directed towards sons on males’ rank post-dispersal and the mediating effects of males’ social behaviour post-dispersal (N = 38). Prox. = proximity, groom. = grooming. A paths represent effects of predictors on males’ social network metrics; b paths represent effects of social network metrics on male rank post-dispersal; ć paths are the total effects of the predictor on male rank; indirect paths are calculated as a x b. Sample sizes reflect the intersection of males with complete data for that specific predictor– mediator–outcome combination.

| Mediator | Path | Mean | 95% CI | pd |
|---|---|---|---|---|
| prox_strength | a | 0.322 | [−0.137, 0.811] | 91.6% |
| prox_strength | b | −0.085 | [−0.440, 0.292] | 68.9% |
| Mediator | Path | Mean | 95% CI | pd |
| prox_strength | c′ | −0.002 | [−0.362, 0.364] | 50.7% |
| prox_strength | indirect | −0.030 | [−0.207, 0.108] | 66.4% |
| prox_degree | a | 0.024 | [−0.323, 0.379] | 55.2% |
| prox_degree | b | −0.096 | [−0.446, 0.261] | 71.0% |
| prox_degree | c′ | −0.032 | [−0.377, 0.325] | 57.7% |
| prox_degree | indirect | −0.002 | [−0.085, 0.073] | 52.2% |
| groom_out_strength | a | 0.135 | [−0.206, 0.478] | 78.6% |
| groom_out_strength | b | −0.260 | [−0.621, 0.109] | 92.0% |
| groom_out_strength | c′ | −0.011 | [−0.341, 0.326] | 52.9% |
| groom_out_strength | indirect | −0.035 | [−0.182, 0.066] | 74.1% |
| groom_in_strength | a | 0.184 | [−0.162, 0.535] | 85.6% |
| groom_in_strength | b | −0.071 | [−0.440, 0.297] | 65.0% |
| groom_in_strength | c′ | −0.019 | [−0.370, 0.340] | 54.6% |
| groom_in_strength | indirect | −0.013 | [−0.131, 0.079] | 60.7% |
| groom_out_degree | a | −0.004 | [−0.338, 0.333] | 51.1% |
| groom_out_degree | b | −0.328 | [−0.667, 0.023] | 96.7% |
| groom_out_degree | c′ | −0.052 | [−0.372, 0.276] | 63.1% |
| groom_out_degree | indirect | 0.001 | [−0.132, 0.135] | 51.0% |
| groom_in_degree | a | −0.035 | [−0.372, 0.306] | 58.6% |
| groom_in_degree | b | −0.118 | [−0.467, 0.234] | 75.0% |
| groom_in_degree | c′ | −0.038 | [−0.383, 0.319] | 59.1% |
| groom_in_degree | indirect | 0.004 | [−0.072, 0.090] | 54.2% |

**Table S2:** Posterior mean, 95% credible interval (equal-tailed quantile), and posterior probability of direction (pd = the larger of prop_positive/prop_negative) for every path in the model testing the effect of maternal proximity to sons on males’ rank post-dispersal and the mediating effects of males’ social behaviour post-dispersal (N = 38). Prox. = proximity, groom. = grooming. A paths represent effects of predictors on males’ social network metrics; b paths represent effects of social network metrics on male rank post-dispersal; ć paths are the total effects of the predictor on male rank; indirect paths are calculated as a x b. Sample sizes reflect the intersection of males with complete data for that specific predictor–mediator–outcome combination.

| Mediator | Path | Mean | 95% CI | pd |
|---|---|---|---|---|
| prox_strength | a | −0.296 | [−0.760, 0.163] | 90.1% |
| prox_strength | b | −0.047 | [−0.374, 0.301] | 62.0% |
| prox_strength | c′ | 0.270 | [−0.084, 0.624] | 93.5% |
| prox_strength | indirect | 0.014 | [−0.116, 0.160] | 59.6% |
| prox_degree | a | −0.127 | [−0.463, 0.205] | 77.6% |
| prox_degree | b | −0.075 | [−0.406, 0.260] | 67.6% |
| prox_degree | c′ | 0.277 | [−0.068, 0.621] | 94.4% |
| prox_degree | indirect | 0.009 | [−0.061, 0.101] | 59.8% |
| groom_out_strength | a | −0.251 | [−0.592, 0.090] | 92.8% |
| groom_out_strength | b | −0.222 | [−0.569, 0.130] | 89.6% |
| groom_out_strength | c′ | 0.248 | [−0.091, 0.586] | 92.7% |
| groom_out_strength | indirect | 0.056 | [−0.044, 0.216] | 83.9% |
| groom_in_strength | a | −0.241 | [−0.593, 0.111] | 91.3% |
| groom_in_strength | b | −0.036 | [−0.389, 0.318] | 58.2% |
| groom_in_strength | c′ | 0.279 | [−0.070, 0.629] | 94.3% |
| groom_in_strength | indirect | 0.009 | [−0.101, 0.131] | 56.8% |
| groom_out_degree | a | −0.199 | [−0.530, 0.134] | 88.4% |
| groom_out_degree | b | −0.297 | [−0.624, 0.037] | 96.1% |
| groom_out_degree | c′ | 0.253 | [−0.076, 0.579] | 93.6% |
| groom_out_degree | indirect | 0.059 | [−0.044, 0.217] | 85.3% |
| groom_in_degree | a | −0.139 | [−0.474, 0.198] | 79.6% |
| groom_in_degree | b | −0.097 | [−0.431, 0.241] | 71.9% |
| groom_in_degree | c′ | 0.278 | [−0.067, 0.621] | 94.4% |
| groom_in_degree | indirect | 0.013 | [−0.060, 0.113] | 63.0% |

**Table S3:** Posterior mean, 95% credible interval (equal-tailed quantile), and posterior probability of direction (pd = the larger of prop_positive/prop_negative) for every path in the model testing the effect of maternal rank on males’ rank post-dispersal and the mediating effects of males’ social behaviour post-dispersal (N = 33). Prox. = proximity, groom. = grooming. A paths represent effects of predictors on males’ social network metrics; b paths represent effects of social network metrics on male rank post-dispersal; ć paths are the total effects of the predictor on male rank; indirect paths are calculated as a x b. Sample sizes reflect the intersection of males with complete data for that specific predictor–mediator–outcome combination. Bold marks the path which 95% credible interval excludes 0.

| Mediator | Path | Mean | 95% CI | pd |
|---|---|---|---|---|
| prox_strength | a | 0.125 | [−0.376, 0.654] | 68.4% |
| prox_strength | b | −0.079 | [−0.442, 0.306] | 67.7% |
| prox_strength | c′ | −0.131 | [−0.502, 0.240] | 76.2% |
| prox_strength | indirect | −0.010 | [−0.142, 0.099] | 56.8% |
| prox_degree | a | −0.286 | [−0.642, 0.068] | 94.5% |
| prox_degree | b | −0.312 | [−0.685, 0.074] | 94.6% |
| prox_degree | c′ | −0.232 | [−0.595, 0.134] | 89.9% |
| prox_degree | indirect | 0.089 | [−0.039, 0.294] | 89.7% |
| groom_out_strength | a | 0.018 | [−0.362, 0.399] | 53.9% |
| groom_out_strength | b | −0.335 | [−0.701, 0.042] | 96.1% |
| groom_out_strength | c′ | −0.142 | [−0.487, 0.203] | 79.7% |
| groom_out_strength | indirect | −0.006 | [−0.167, 0.146] | 53.7% |
| groom_in_strength | a | 0.193 | [−0.195, 0.585] | 83.9% |
| groom_in_strength | b | −0.158 | [−0.549, 0.243] | 79.0% |
| groom_in_strength | c′ | −0.116 | [−0.487, 0.254] | 73.7% |
| groom_in_strength | indirect | −0.031 | [−0.188, 0.074] | 69.8% |
| groom_out_degree | a | 0.042 | [−0.323, 0.408] | 59.0% |
| groom_out_degree | b | **−0.429** | **[−0.771, −0.076]** | **99.0%** |
| groom_out_degree | c′ | −0.135 | [−0.465, 0.195] | 79.7% |
| groom_out_degree | indirect | −0.018 | [−0.203, 0.155] | 58.9% |
| groom_in_degree | a | 0.094 | [−0.282, 0.477] | 68.9% |
| groom_in_degree | b | −0.220 | [−0.589, 0.155] | 88.1% |
| groom_in_degree | c′ | −0.131 | [−0.490, 0.230] | 76.8% |
| groom_in_degree | indirect | −0.021 | [−0.161, 0.085] | 64.4% |

**Table S4:** Posterior mean, 95% credible interval (equal-tailed quantile), and posterior probability of direction (pd = the larger of prop_positive/prop_negative) for every path in the model testing the effect of maternal centrality in grooming networks on males’ rank post-dispersal and the mediating effects of males’ social behaviour post-dispersal (N = 38). Prox. = proximity, groom. = grooming. A paths represent effects of predictors on males’ social network metrics; b paths represent effects of social network metrics on male rank post-dispersal; ć paths are the total effects of the predictor on male rank; indirect paths are calculated as a x b. Sample sizes reflect the intersection of males with complete data for that specific predictor– mediator–outcome combination.

| **Mediator** | **Path** | **Mean** | **95% CI** | **pd** |
|---|---|---|---|---|
| prox_strength | a | 0.350 | [−0.096, 0.840] | 93.7% |
| prox_strength | b | −0.098 | [−0.453, 0.293] | 71.5% |
| prox_strength | c′ | 0.049 | [−0.312, 0.410] | 60.8% |
| prox_strength | indirect | −0.034 | [−0.216, 0.116] | 68.8% |
| prox_degree | a | −0.253 | [−0.579, 0.076] | 93.6% |
| prox_degree | b | −0.099 | [−0.465, 0.274] | 70.6% |
| prox_degree | c′ | −0.008 | [−0.365, 0.350] | 51.9% |
| prox_degree | indirect | 0.025 | [−0.084, 0.164] | 68.0% |
| groom_out_strength | a | 0.137 | [−0.211, 0.489] | 78.2% |
| groom_out_strength | b | −0.269 | [−0.629, 0.099] | 92.8% |
| groom_out_strength | c′ | 0.048 | [−0.283, 0.382] | 61.4% |
| groom_out_strength | indirect | −0.037 | [−0.190, 0.068] | 74.1% |
| groom_in_strength | a | 0.278 | [−0.082, 0.654] | 93.6% |
| groom_in_strength | b | −0.087 | [−0.465, 0.297] | 67.8% |
| groom_in_strength | c′ | 0.040 | [−0.317, 0.398] | 59.0% |
| groom_in_strength | indirect | −0.024 | [−0.181, 0.101] | 65.6% |
| groom_out_degree | a | −0.072 | [−0.414, 0.283] | 66.4% |
| groom_out_degree | b | −0.329 | [−0.672, 0.023] | 96.7% |
| groom_out_degree | c′ | −0.016 | [−0.340, 0.313] | 54.2% |
| groom_out_degree | indirect | 0.024 | [−0.107, 0.174] | 65.3% |
| groom_in_degree | a | −0.024 | [−0.372, 0.339] | 56.1% |
| groom_in_degree | b | −0.117 | [−0.466, 0.235] | 74.8% |
| groom_in_degree | c′ | 0.015 | [−0.326, 0.357] | 53.3% |
| groom_in_degree | indirect | 0.003 | [−0.078, 0.089] | 53.0% |

**Table S5:** Posterior mean, 95% credible interval (equal-tailed quantile), and posterior probability of direction (pd = the larger of prop_positive/prop_negative) for every path in the model testing the effect of maternal centrality in proximity networks on males’ rank post-dispersal and the mediating effects of males’ social behaviour post-dispersal (N = 38). Prox. = proximity, groom. = grooming. A paths represent effects of predictors on males’ social network metrics; b paths represent effects of social network metrics on male rank post-dispersal; ć paths are the total effects of the predictor on male rank; indirect paths are calculated as a x b. Sample sizes reflect the intersection of males with complete data for that specific predictor– mediator–outcome combination.

| **Mediator** | **Path** | **Mean** | **95% CI** | **pd** |
|---|---|---|---|---|
| prox_strength | a | −0.155 | [−0.631, 0.313] | 74.6% |
| prox_strength | b | −0.084 | [−0.426, 0.283] | 69.4% |
| prox_strength | c′ | 0.039 | [−0.306, 0.380] | 59.2% |
| prox_strength | indirect | 0.012 | [−0.090, 0.138] | 59.4% |
| prox_degree | a | −0.133 | [−0.468, 0.202] | 78.7% |
| prox_degree | b | −0.092 | [−0.442, 0.265] | 70.1% |
| prox_degree | c′ | 0.041 | [−0.307, 0.382] | 59.6% |
| prox_degree | indirect | 0.012 | [−0.064, 0.113] | 61.5% |
| groom_out_strength | a | −0.193 | [−0.540, 0.157] | 86.6% |
| groom_out_strength | b | −0.261 | [−0.625, 0.110] | 92.0% |
| groom_out_strength | c′ | 0.013 | [−0.325, 0.346] | 53.2% |
| groom_out_strength | indirect | 0.050 | [−0.053, 0.214] | 80.7% |
| groom_in_strength | a | −0.187 | [−0.540, 0.173] | 85.4% |
| groom_in_strength | b | −0.071 | [−0.440, 0.298] | 64.8% |
| groom_in_strength | c′ | 0.038 | [−0.312, 0.383] | 58.8% |
| groom_in_strength | indirect | 0.013 | [−0.083, 0.132] | 60.3% |
| groom_out_degree | a | −0.094 | [−0.436, 0.242] | 70.9% |
| groom_out_degree | b | −0.325 | [−0.665, 0.025] | 96.6% |
| groom_out_degree | c′ | 0.037 | [−0.290, 0.358] | 59.4% |
| groom_out_degree | indirect | 0.030 | [−0.090, 0.182] | 69.5% |
| groom_in_degree | a | −0.101 | [−0.438, 0.233] | 72.6% |
| groom_in_degree | b | −0.113 | [−0.462, 0.239] | 74.1% |
| groom_in_degree | c′ | 0.040 | [−0.305, 0.380] | 59.5% |
| groom_in_degree | indirect | 0.011 | [−0.061, 0.110] | 60.9% |

